# Super-resolved live imaging of thick biological samples with 3D Random Illumination Microscopy (3D-RIM)

**DOI:** 10.1101/2025.08.06.668879

**Authors:** Thomas Mangeat, Lorry Mazzella, Benoît Rogez, Guillaume Giroussens, Xiangyi Li, Mathilde Bernard, Pablo Vargas, Marc Allain, Simon Labouesse, Jérôme Idier, Loïc LeGoff, Anne Sentenac

## Abstract

Super-resolved volume imaging of thick, live specimens is greatly hampered by sample induced aberrations and out-of-focus fluorescence. In this work, using a combination of speckled illuminations, remote focusing, three-dimensional photon reassignement and variance processing, we obtained volume images with super-resolution (110 nm transverse, 270 nm axial) on thin samples and maintained a high contrast and high resolution level throughout tens of microns of highly aberrant tissues and up to hundreds of microns in collagen scaffolds.

---

Super-resolved live imaging of thick biological samples, spanning several tens of micrometers, is becoming increasingly important in cell and developmental biology, but remains a major challenge for one-photon fluorescence microscopy [1, 2]. All techniques are affected by sample-induced aberrations on the detection side. In addition, methods employing uniform illumination suffer from strong out-of-focus fluorescence background, whereas techniques using patterned illuminations (such as confocal-related methods, periodic Structured Illumination Microscopy (SIM), and lattice light-sheet microscopy) are impacted by sample-induced scattering and aberrations on the excitation side, which further degrade their resolution as the imaging depth increases. In this work, we present three-dimensional (3D) random illumination microscopy (3D-RIM) a super-resolved widefield microscopy method adapted to volume imaging that is insensitive to aberrations on the excitation side and efficiently removes the out-of-focus fluorescence.

3D-RIM forms a super-resolved volume image of the sample from the variance of multiple low-resolution images recorded under random speckled illuminations [3, 4, 5, 6]. The key feature of 3D-RIM lays in a global 3D data processing that requires 3D low-resolution images, 3D photon reassignement and 3D variance matching. In brief, we take advantage of the information brought about by the out-of-focus photons and exploit the 3D properties of the point spread function and speckle statistics. As the sample should not be moved with respect to the speckled illumination when recording the 3D low-resolution image, a remote focusing unit was introduced in the microscope’s detection arm to perform axial scanning, (see Fig. 1a and supplementary methods for details on the experimental set-up and 3D data processing).

**Figure 1.**
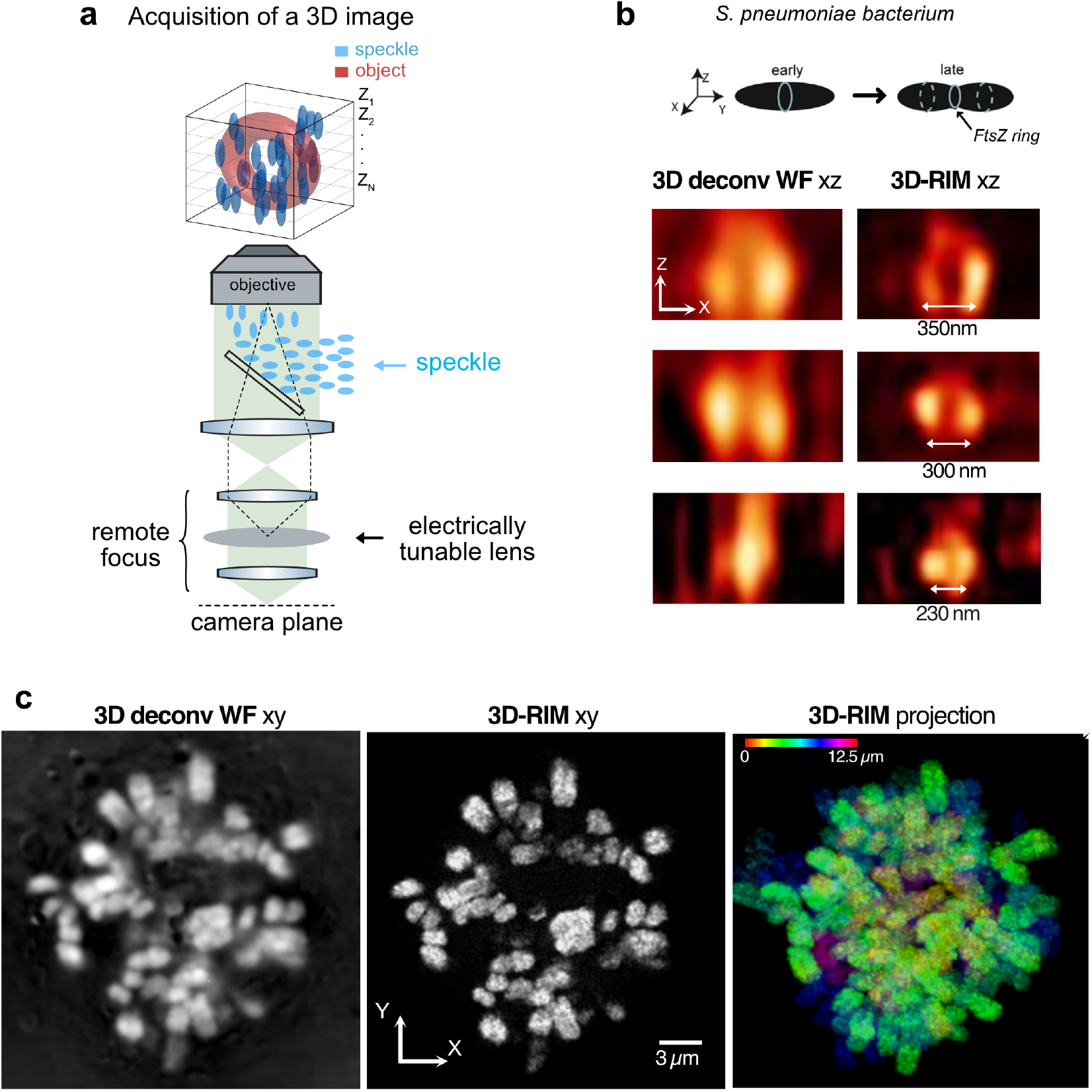
3D-RIM in single cells. **a**: Schematic of the 3D-RIM set-up. The sample (here a FtsZ ring) is excited by a speckled illumination. A remote focusing unit, using an electrically tunable lens, permits to record z-stack images while keeping the illumination of the sample unchanged. Multiple (100 to 200) low-resolution 3D speckled images of a sample are recorded under different random speckled illuminations. The super-resolved 3D image is obtained from the variance of the speckled images using a variance matching algorithm (see the methods). **b**: FtsZ:GFP fusion in a live Streptococcus pneumonia bacterium during cell division (the excitation and emission wavelengths are *λ*_exc_ = 488 nm and *λ*_em_ = 522 nm respectively, the Numerical Aperture of the objective is NA = 1.49). Schematic: FtsZ assembles into a ring that gradually shrinks as cytokinesis proceeds. XZ cuts of 3D images of closing rings at three different stages, obtained with 3D-RIM or deconvolved widefield microscopy. In sharp contrast with the deconvolved widefield images, the ring structure is well seen in the 3D-RIM images and the internal hole can even be suspected when the ring structure shrinks to a total diameter of 230 nm. **c**: A fixed human Hep3B cell stained for DNA (*λ*_exc_ = 405 nm, *λ*_em_ = 450 nm, NA = 1.3).The sample is 15 microns thick. From left to right : one XY cut of the deconvolved widefield 3D image, the same cut imaged by 3D-RIM, the maximum intensity projection of the 3D-RIM image with colors encoding Z, see movie 1 for a 3D representation. The 3D-RIM resolution estimated by FRC is 110 nm transversally and 270 nm axially.

We first demonstrate the performance of 3D-RIM on thin biological samples. We imaged FtsZ:GFP fusions in live Streptococcus pneumonia bacteria. FtsZ labelling enables the evolution of the construction ring to be tracked throughout the division process. Figure 1b shows vertical cuts (XZ) of such rings at different stages of cell division. 3D-RIM could visualize well the central hole of vertical rings with an overall diameter close to 300 nm. Figure 1c shows the chromosomes of fixed human cells labelled for DNA. 3D-RIM reconstructions exhibited a better resolution, notably along the optical axis, better contrast, and less noise than 3D-deconvolved widefield images. The rightmost panel in Fig. 1c displays a maximum intensity projection from 3D-RIM, with color encoding position along the optical axis. A 3D representation of the volume image is provided in the supplementary movie 1. The Fourier Ring Correlation (FRC) analysis [7] estimated the transverse and axial resolution to 110 nm and 270 nm, respectively. Thus, on thin samples, 3D-RIM matched the resolution of 3D-SIM [8, 9].

We next explored the ability of 3D-RIM to image thick biological tissues. We considered *Drosophila* embryos (stage before invagination) that had been immunostained for lamin in order to outline the nuclear envelopes. We formed a 3D-RIM volume image of the embryo, ranging from 27 to 52 *µ*m in depth. Figure 2a displays a large-scale XY cut of the 3D image at a depth of 40 *µ*m from the coverslip giving a global view of the embryo circumference. The XZ and XY cuts throughout the volume are shown in the supplementary movies 2 and 3. The inset in Fig. 2a provides a maximum intensity projection with color encoding depth, offering a comprehensive view of the nuclei architecture. The transverse and axial resolutions were estimated by FRC to about 180 nm and 450 nm, respectively. For comparison, we also formed a 3D image by translating the sample through the focal plane and stacking the super-resolved, optically sectioned, two-dimensional (2D) reconstructions of regular RIM [3]. As shown in Fig. 2b, in the zoomed insets and the accompanying line profiles along the optical axis (z), 3D-RIM provided a better image contrast and resolution. In Fig. 2c,d, we imaged the Paneth cells of a mouse intestine that had been stained with Wheat Germ Agglutinin (WGA) conjugated to rhodamine to highlight the membranes of secretory granules. High-resolution imaging in this tissue is particularly challenging due to the dense fluorescence labeling and the optically scattering and aberrating environment created by the secretory granules and the complex shape of the intestin crypts. We formed a super-resolved 3D-RIM volume image of the tissue from 27 *µ*m to 42 *µ*m of depth from the coverslip. The maximum intensity projection, color-coded for depths, revealed intricate details of the tissue architecture (Fig. 2c). Figure 2c,d displays XZ cuts of the 3D image and two intensity profiles along the z-axis comparing 3D-RIM and regular RIM. For this densely labeled sample, regular RIM was hampered by the noisy out-of-focus signal in the raw images. The stack of 2D reconstructions was marred by a strong background, making the individual granules hard to distinguish. In contrast, the absence of background due to efficient noise removal and optical sectioning allowed 3D-RIM to delineate the precise shape of individual granules successfully. The transverse and axial resolutions of 3D-RIM were estimated by FRC to 230 nm and 470 nm, respectively.

**Figure 2.**
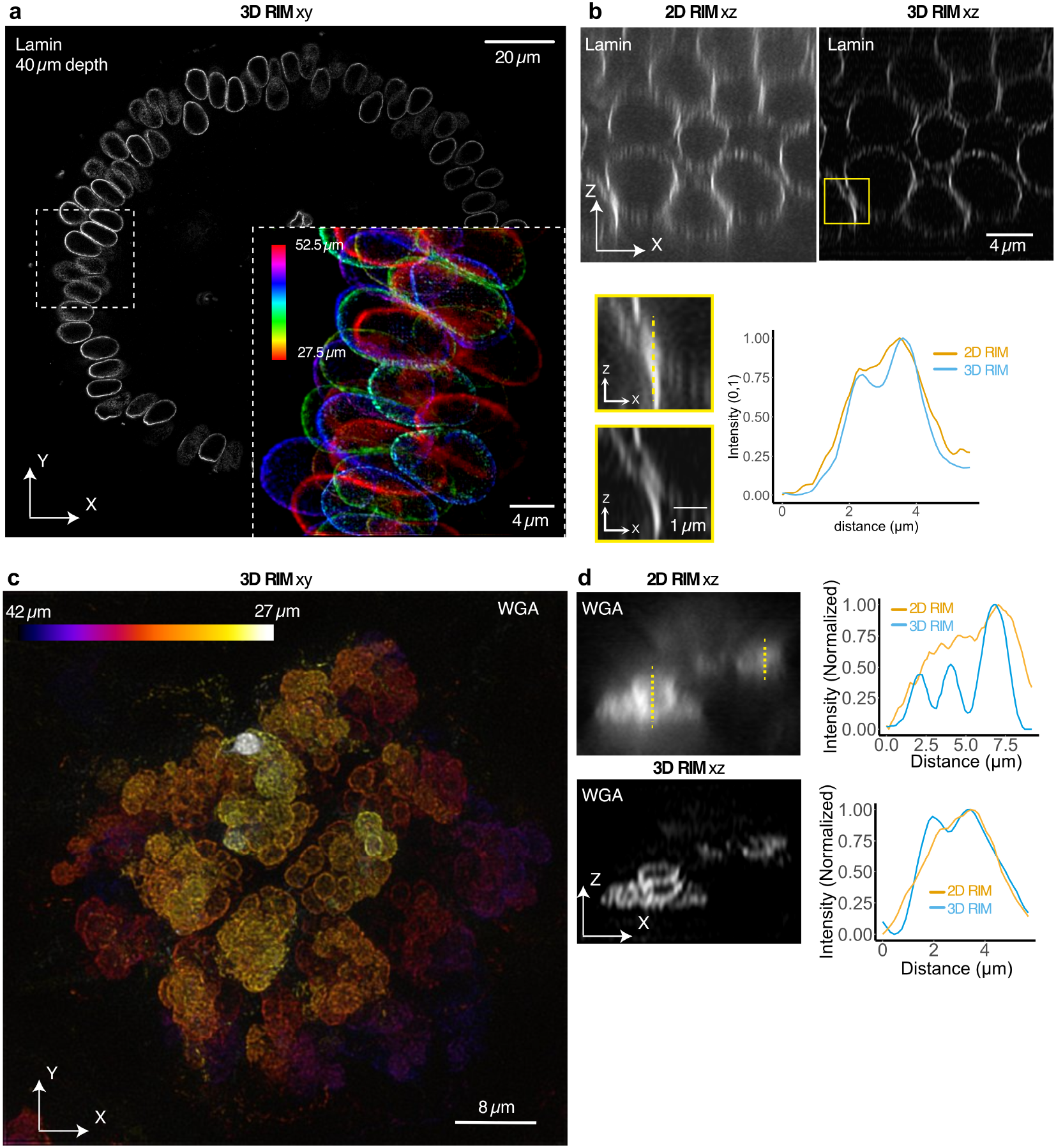
3D-RIM in thick biological tissues. **a**,**b**: *Drosophila* embryo with lamin immunolabeling targeting the nuclear envelopes (*λ*_exc_ = 488 nm, *λ*_em_ = 522 nm, NA = 1.3). **a**: large-scale XY cut of the 3D-RIM image taken 40 *µ*m deep from the coverslip. The many nuclei accumulate at the surface of the embryo (see also movies 2,3). The inset shows the maximum intensity projection (colors coding for Z) of a 3D-RIM image spanning from 27 *µ*m to 52 *µ*m from the coverslip. The quality of the reconstruction is maintained throughout the 25 *µm* of axial scan. The transverse and axial resolutions are estimated by FRC to 180 nm and 450 nm, respectively. **b**: XZ cuts of the 3D images of the same sample obtained with 3D-RIM or regular RIM (2D-RIM). The inset provides zooms on the selected region of interest. The intensity profiles displays the increased resolution of 3D-RIM along the optical axis (Z) (blue line) as compared to regular RIM (orange). **c**,**d**: Paneth cells of a mouse intestine in which the saccharides are stained with WGA conjugated to rhodamin (*λ*_exc_ = 561 nm, *λ*_em_ = 596 nm, NA = 1.3). **c**: Maximum intensity projection of a 3D-RIM image spanning from 27 *µ*m to 42 *µ*m from the coverslip. The transverse and axial resolutions are estimated by FRC to 230 nm and 470 nm, respectively. **d**: Representative XZ cut of the 3D image comparing 3D-RIM with regular RIM (2D RIM). Two intensity profiles along Z are displayed. 3D-RIM clearly outlined the shape of the granules (blue line). Regular RIM was plagued by strong out-of-focus noise (orange line).

The robustness of 3D-RIM to aberrations and its efficient out-of-focus fluorescence removal is further evidenced in supplementary movies 4,5, and 6 which show 3D representations of the tubulin in dendritic cells migrating in a collagen gel scaffold. The three volume images taken at 40 *µm*, 150 *µ*m and 450 *µ*m from the coverslip are well contrasted and, surprisingly, have similar axial resolutions, from 750 nm to 1 *µ*m and same transverse resolutions, about 330 nm. In contrast, the 3D deconvolved widefield image suffers from a strong out-of-focus background, and from a twice lower resolution (see supplementary Fig. 1).

Finally, we illustrate in supplementary figure 2 and movies 7, 8, and 9 the ability of 3D-RIM to provide dynamic volume images of live thick specimen. We imaged fixed and live border cells during their migration in the Drosophila egg chamber. Individual actin fiber bundles were remarkably well resolved in fixed border cells 33 to 46 *µ*m from the coverslip (Fig. S2a-c), and, in live egg chambers, actin dynamic protrusions could be followed during the whole migration process (lasting about 30 minutes and reaching a depth of 65 *µ*m) without bleaching nor apparent phototoxicity (Fig. S2d and movie 9).

In conclusion, by combining the efficiency of a global 3D data processing with the robustness to aberrations, low phototoxicity and ease of use of RIM [3] on living samples, 3D-RIM provides well-contrasted volume images of thick biological samples at resolutions and depths never documented before. This technique opens up new possibilities for investigating subcellular structures and dynamics in intact tissues like developing embryos or organoids.

## Acknowledgments

This work was funded by the following agencies: Agence Nationale de la Recherche (ANR-20-CE45-0024, ANR-22-CE13-0039, ANR-22-CE42-0010, ANR-22-CE42-0026, ANR-24-CE13-2163); Institut Carnot star (3D-RIM).

This project is funded by the *≪* France 2030 *≫* investment plan managed by the French National Research Agency (IDEC, ANR-21-ESRE-0002), and from Excellence Initiative of Aix-Marseille University - A*MIDEX.

We would like to thank Nathalie Campo and Isabelle Mortier from the Polar team at LMGM CBI for producing the S. pneumoniae cultures and strains and Maëlle Carraz from the Bystricky team at MCD CBI for culturing, fixing and labelling the Hep3B human hepatocarcinoma cells shown in Figure 1. Our thanks also go to Justine Creff from the Krndija team at MCD CBI, who prepared the mouse intestine section samples and performed the immunostaining and to Ronan Bouzignac from the Suzanne team at MCD CBI, who prepared the Drosophila embryo sections and performed the immunostaining used in Figure 2. We thank Léane Lesbats for help with 3D-IF of dendritic cells in Figure 1S. Finally, we would like to thank Halo Li and the Wang team at MCD CBI for providing the fixed and live Drosophila ovarian chamber samples shown in Figure 2S.

## Methods

### 3D-RIM data processing

3D-RIM involves the recording of multiple 3D low-resolution images of the sample under different random speckled illuminations. The key requirement of 3D-RIM data processing is that each 3D speckled image *I* be obtained by keeping the speckled illumination of the sample unchanged so that it can be modeled as the convolution between a 3D point spread function *h* and the product of the sample 3D fluorescence density *ρ* and the 3D speckle intensity pattern *S*,

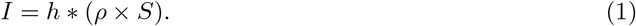

The reconstruction method, 3DalgoRIM, proceeds by first deconvolving the raw images using a 3D Wiener filter, *I*^*′*^ = *I f* with 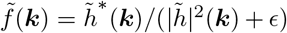, where *ã* stands for the 3D Fourier transform of *a, b*^*∗*^ is the conjugate of *b*, and *ϵ* is a regularization parameter that is tuned manually. Then, 3DalgoRIM calculates the empiric variance of the deconvolved speckled images 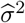. Note that the 3D deconvolution of the raw 3D images is a crucial step. Reassigning the out-of-focus photons to their sources amplifies the peaks of fluorescence intensity stemming from the speckle hot spots and improves dramatically the signal to noise ratio of the images’ variance.

Last, 3DalgoRIM estimates the 3D fluorescence density *ρ* iteratively so as to minimize the distance between the empirical standard deviation and its theoretical model,

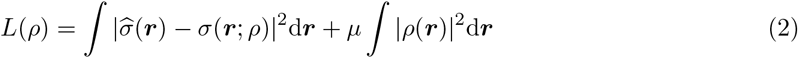

with *µ* a Tikhonov parameter and *σ*(***r***; *ρ*) the theoretical standard deviation derived by assuming an infinite number of speckle realizations,

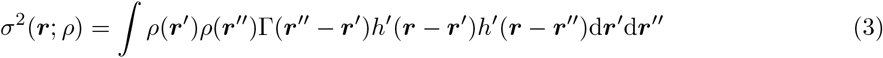

with *h*^*′*^ = *h* ∗ *f*, Γ the 3D speckle auto-covariance, Γ(***r***−***r***^*′*^) = ⟨(*S*(***r***) − ⟨*S*⟩)(*S*(***r***^*′*^) − ⟨*S*⟩)⟩ with ⟨⟩ is the averaging over the speckle realizations. Note that the calculation of the 3D theoretical standard deviation is made tractable (and relatively fast) thanks to a truncated spectral decomposition of the positive definite operator *T* defined by *T* (***u, v***) = *h*^*′*^(***u***)*h*^*′*^(***v***)Γ(***u*** − ***v***),

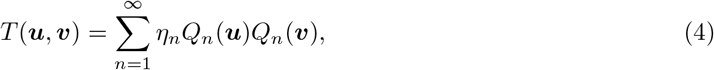

where *η*_*n*_ are positive decreasing values. In all the configurations we studied, the first ten terms of the sum were sufficient to get an accurate estimation of *σ*. The minimization of Eq. (2) is performed with a preconditioned conjugate gradient algorithm that is described in detail in [10]. It is worth noting that our reconstruction technique uses only a standard Tikhonov regularization.

The 2D reconstructions of regular RIM were also performed by 3DalgoRIM except that each plane was processed independently by setting the dimension along the optical axis of the discretized 3D functions *h*, Γ and *ρ* to one.

In all the reconstructions, the microscope point spread function *h* and the speckle auto-covariance Γ were estimated using the point spread function generator of Image J (the Gibson-Lanni model) [11] at the illumination or emission wavelengths. The tuning of the regularization parameters *ϵ* and *µ*, which depends on the signal to noise ratio of the raw images, is done manually. For all the considered samples, *ϵ* varied between 5 10^*−*4^ and 2 10^*−*3^, and *µ* between 1.6 10^*−*7^ and 6 10^*−*6^. The iterative process converged in 50 to 100 iterations.

### Experimental set-up

The 3D-RIM was constructed using a standard Nikon TiE fluorescence microscope in an epi configuration. Light from diode and solid-state lasers (405 nm, 488 nm, 561 nm and 633 nm) was passed into a binary Spatial Light Modulator (SLM, 4D QXGA), which was conjugated to the image plane of the objective via an intermediate lens (AC254-250-A-ML, Thorlabs, f = 250 mm), to generate speckles.

The SLM was set up for phase modulation, following [12], using a polarisation beam splitter and half-wave plate. In this configuration, the zero order of the SLM is suppressed. The objectives used were: i-a 60x silicon oil objective (Nikon, PLAN APO 60X/1.30 SIL NA=1.3) for Hep3B cells, *Drosophila* embryos, Paneth cells of the mouse intestine and border cells in *Drosophila* egg chambers; ii-a 100x oil immersion (Nikon, 100x 1.49 APO SR TIRF) for FtsZ rings; iii-a 40x water immersion (CFI Apo LWD Lambda S 40XC WI, NA=1.15) for the dendritic cells in collagen gel scaffolds.

In most microscopes, the z-scan of the sample is performed by translating the sample (or equivalently the objective) through the focal plane. Now, if the illumination is inhomogeneous along the optical axis, as with speckled illumination, translating the sample modifies the way the sample is illuminated. Then, the image model is not a 3D convolution anymore, which prevents the full application of 3D-RIM data processing. Hence, the key feature of 3D-RIM set-up lays in the remote focusing unit that permits to produce 3D-stack of images without moving the sample with respect to the illumination. The remote focusing unit consisted of a telecentric 4f relay using two lenses of focal 75mm (AC254-075-A-ML Thorlabs) and 100mm (AC254-100-A-ML Thorlabs) respectively to reimage the intermediate image plane on the camera (Kinetix Scientific Teledyne) and a gravity corrected electrically tunable lens placed in the Fourier plane of the 4f relay (Optotune, EL-16-40-GTC-VIS-5D-1-C). The optical power of the electrically tunable lens was controlled through an analog signal sent to its controller (Optotune, ICC-1C Controller KIT).

As changing the z-focus is a limiting factor in the temporal sequence, we recorded P speckled images at a given plane, followed by a z-shift and re-imaging of the same speckle sequence at the new z plane (P=200 on fixed samples and P=100 on live samples). A z-step of 150 nm was used for the imaging of *Drosophila* embryos, Paneth cells of the mouse intestine, and fixed border cells. A step of 90 nm was used for the FtsZ rings and Hep3B chromosomes and a step of 250 nm was used for the live imaging of border cells and the imaging of dendritic cells.

The typical exposure time for one speckled image plane was 10 ms. Synchronisation and acquisition were performed by the commercial Inscoper L electronic/software system.

## Samples preparation

### FtsZ rings of S. pneumoniae

S. pneumoniae strain containing the FtsZ-GFP fusion (R3702, Bergé et al., 2013), was grown in C+Y medium as previously described at 37°C to an OD550 of 0.15 (Bergé et al., 2017). Samples were collected, pelleted (3 min, 3,000 g) and resuspended in cold C medium. The C medium contained per liter: 5 g casein hydrolysate, 6 mg tryptophane, 11.25 mg cysteine, 2 g sodium acetate and 8.5 g K2HPO4. Cells were spotted on a microscope slide containing a slab of 1.2% agarose in C medium and covered with a pre-treated coverslip before imaging.

### Hep3B cell DNA staining

Human hepatocarcinoma cells Hep3B (ATCC HB-8064) were seeded in *µ*-Dish 35 mm, high Grid-500 (ibidi^®^, Cat 81166) at 250,000 cells per dish overnight in complete DMEM medium supplemented with 10% of SVF. Cells were fixed with 4% paraformaldehyde in PBS for 10 min at room temperature, extensively washed with PBS before their labeling with DAPI (2 *µ*g/mL) for 10 min in the dark, extensively washed, and imaged.

### *Drosophila* embryo preparation

We imaged embryos of genotype kuk[EY07696] (Blomington stock #16856), which bear a viable hypomorphic mutation in the gene kugelkern/charleston, responsible for mild nuclear morphology defects. Embryos were collected 2–4 hours after egg laying (AEL), dechorionated in 50% bleach for 2 minutes, fixed in 1 mL of 37% formaldehyde (Sigma-Aldrich, 818708) and 1 mL of heptane for 5 minutes at 4^*o*^*C*, washed twice in PBS containing 0.3% Triton X-100. Selected embryos were cut in their middle using a microscalpel (Micro Knives–Plastic Handle, 10315-12, F.S.T.), incubated at 4^*o*^C with mouse anti-Lamin antibody (1:10, ADL67.10, DSHB), and 4 hours with goat anti-mouse Alexa Fluor 488 (1:100, Interchim). For imaging, embryo cuts were mounted between two coverslips using a 120*µm* deep spacer (Secure-Seal™, Sigma-Aldrich) in Vectashield mounting medium. **Paneth cells of the mouse intestine**. Jejunum from one sacrificed mouse was fixed in 4% paraformaldehyde (Electron Microscopy Sciences), washed in PBS, sectioned at 200 − 300*µm* thickness, permeabilized with 1% Triton X-100 for 1 h. Saccharides were stained with Wheat Germ Agglutinin (WGA) conjugated to rhodamin, washed in PBS and mounted in Aqua-Poly-Mount, (PolySciences). Mice were housed and procedures performed in accordance with European and national regulations (Protocol Authorisation APAFIS #38994-2023062718345638). Mice were bred in the CBI Specific and Opportunistic Pathogen Free (SOPF) animal facility. Mice were maintained on a C57Bl/6N genetic background.

### Dendritic cells in a collagen matrix

Mouse bone marrow-derived dendritic cells (BMDCs) were differentiated and activated with LPS as previously published [13]. Cells were seeded in a collagen gel (2mg/m, type I rat tail collagen, Corning 354249) as described in [14]. The collagen gel was cast in a PDMS-made microfabricated collagen chamber (height=500µm, width=3mm, length=4.5mm). The chemokine CCL19 (80nM, ThermoFischer 250-27B) was added in the cell medium to trigger chemotaxis. After 1 hour of directional migration, cells were fixed with 4%PFA then permeabilized with 0.3% Triton X-100. Cells were stained for alpha-tubulin (anti-alpha-Tubulin Antibody, Thermofisher, MA1-80017 detected using a AlexaFluor647-coupled anti-rat secondary antibody, Thermofisher, A21472).

### *Drosophila* border cells migration

For live imaging, we used the Drosophila stocks slbo::LifeAct–RFP [15] and followed the protocol of Dai and Montell [16]. For fixed imaging conditions, ovaries from Sqh::RLCmyosinII–GFP Drosophila were dissected in Schneider’s medium and fixed in 4% paraformaldehyde in phosphate-buffered saline (PBS) for 20 minutes at room temperature. After fixation, the egg chambers were rinsed three times with PBS containing 0.3% Triton X-100 (PBST). The egg chambers were incubated with Alexa-561-conjugated phalloidin (1:100 dilution; Invitrogen, A34055) for 2 hours at room temperature to achieve F-actin staining, followed by three 20-minute washes with PBST.

## Supplementary Figures

**Figure S1:**
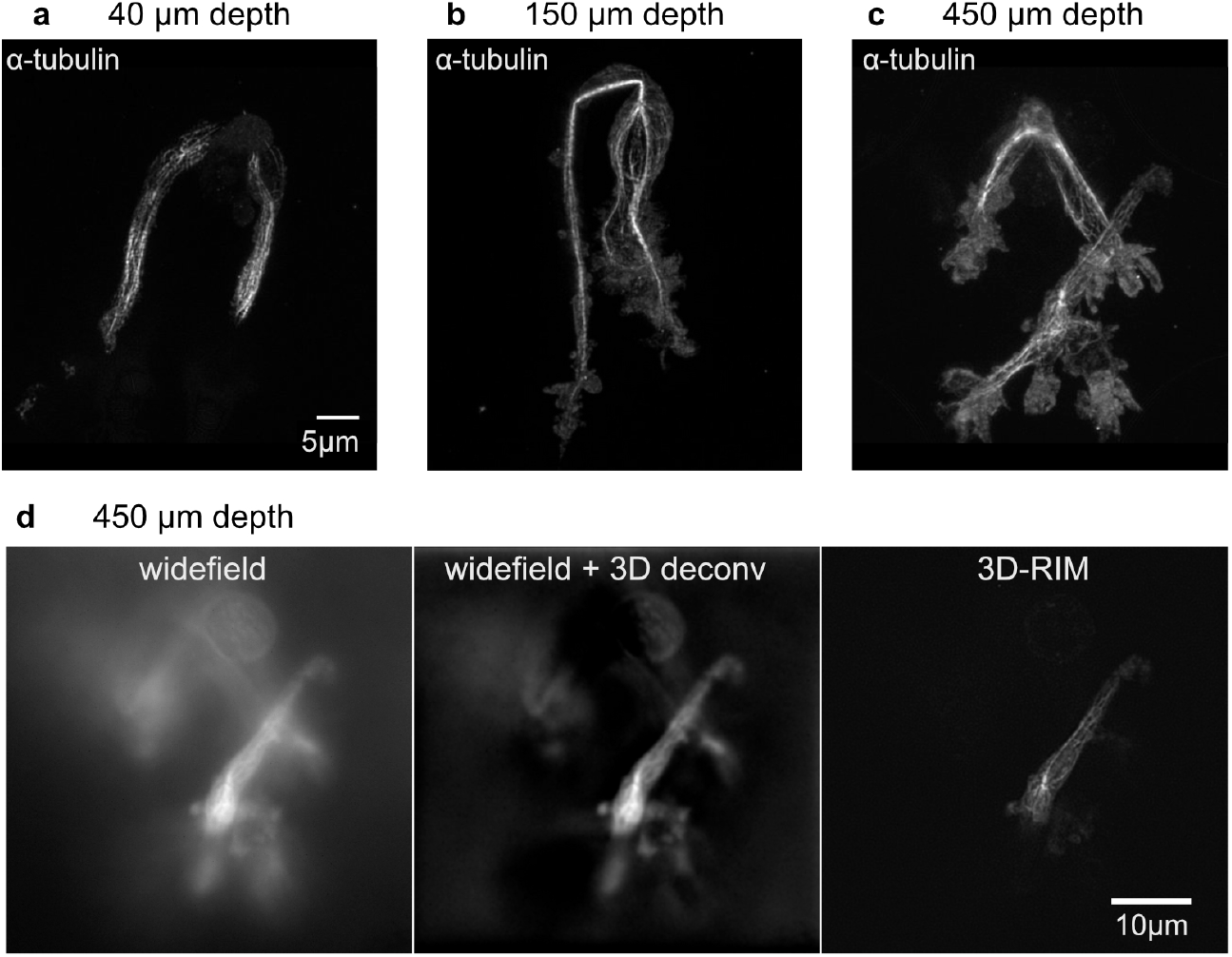
Imaging chemotactic dendritic cells in a collagen scaffold (*λ*_exc_ = 633 nm, *λ*_em_ = 670 nm, NA = 1.15). (a-c)*α*-tubulin immuno-fluorescence in mouse bone marrow-derived dendritic cells imaged at different depths (40*µm*, 150*µm* and 450*µm*). Images are maximum intensity projections (see also movies 4,5,6 for a 3D representation). The resolutions of the three volume images, estimated through analysis of the full width at half maximum of the thinnest filaments, is around 330 nm transversally for all depths and ranging from 750 nm at 40 *µm* to 1000 nm at 150 and 450 *µ*m axially. (d) Comparison of the standard widefield image, 3D-deconvolved widefield image, and 3D-RIM on one section of the cell at 450 *µ*m depth. The transverse resolution of the deconvolved widefield image is estimated around 660 nm.

**Figure S2:**
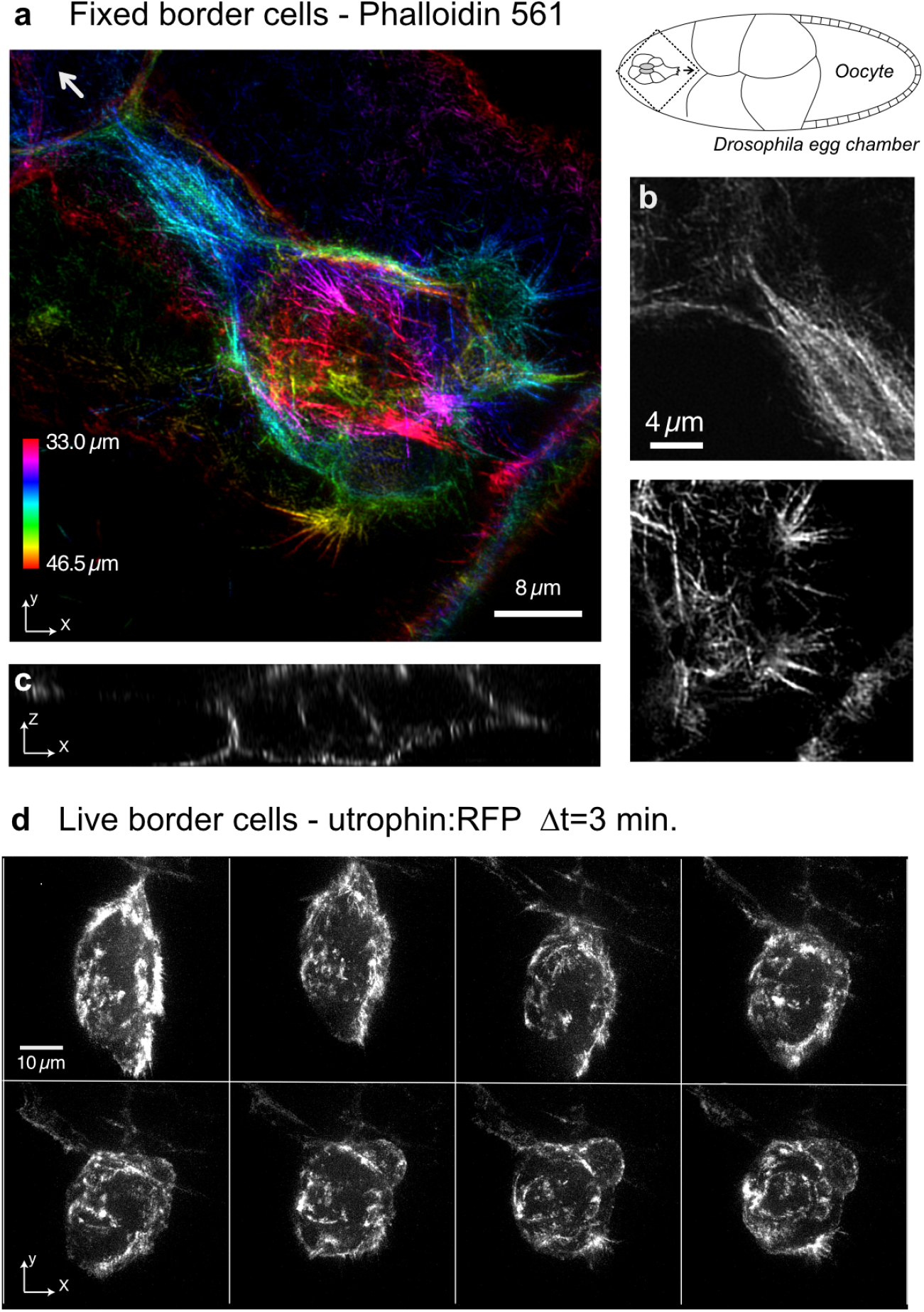
Imaging actin in border cell migration in the *Drosophila* egg chamber. (a-c) Fixed egg chamber with Alexa561 phalloidin staining when the border cells are starting their migration (*λ*_exc_ = 488 nm, *λ*_em_ = 522 nm, NA = 1.3). a: maximum projection with z-encoding colors of the volume image provided in movies 7 and 8. The individual actin bundles are visible throughout the whole border cell cluster, ranging from 33 microns to 46 microns from the coverslip. The schematic on the right shows the structure of the *Drosophila* egg chamber. The dotted square contains the border cells imaged in Fig. S2a. The arrow in the projection and in the schematic shows the direction of migration of border cells, towards the oocyte. b: two XY zooms at different depths and locations. c: one XZ cut of the volume image extracted from movie 7. d: Live 3D imaging of the actin of border cells at the end of their migration when they reach the center of the egg chamber, at a depth of 65 microns from the coverslip, extracted from movie 9 (*λ*_exc_ = 561 nm, *λ*_em_ = 596 nm, NA = 1.3). Z-projection of 8 volume images taken three minutes apart.

## Supplementary movie captions

Movie 1: 3D representation of DNA in a fixed human cell, associated to Fig. 1c.

Movies 2 and 3 : Full stacks of XY and XZ slices of a volume image of lamin in a fixed *Drosophila* embryo, associated to Fig. 2a.

Movies 4,5,6: 3D representation of the tubulin of fixed dendritic cells, 40 *µm*, 150 *µm* and 450 *µm* deep in a collagen scaffold, respectively. The movies are associated to supplementary Fig. S1

Movies 7 and 8 : Full stacks of XY and XZ slices of the volume image of actin in a fixed *Drosophila* egg chamber, associated to supplementary Fig. S2.

Movie 9: Live volume imaging of actin in a *Drosophila* egg chamber during border cells migration. 3D representation of 8 volume images taken 3 mn apart. The movie is associated to supplementary Fig. S2

